# The essential role of disulfide bonds for the hierarchical self-assembly and wet-adhesion of CP20-derived peptides

**DOI:** 10.1101/2022.06.15.496244

**Authors:** Baoshan Li, Junyi Song, Ting Mao, Ling Zeng, Zonghuang Ye, Biru Hu

## Abstract

Barnacles are typical fouling organisms which strongly adhere to immersed solid substrates by secreting proteinaceous adhesives called cement proteins (CPs). The self-assembly of the cement proteins forms a permanently bounded layer that binds barnacle to foreign surfaces. However, due to the abundance of cysteines in whole-length CP20, it is difficult to determine its natural structure and to properly describe its self-assembly properties. In this study, a putative functional motif of *Balanus albicostatus* CP20 (BalCP20) is identified and found to present distinctive self-assembly and wet-adhesion characteristics. The atomic force microscopy (AFM) and transmission electron microscope (TEM) investigations show that wildtype BalCP20-P3 forms grain-like spindles, which further assembly into fractal-like structures looks like ears of wheat. SDS-PAGE, AFM and LSCM show that DTT treatment opens up disulfide bonds between cysteines and disrupts the fractal-like structures (eras of wheat). Additionally, these morphologies are abolished when one of the BalCP20-P3 four cysteines is mutated by alanine. Circular dichroism (CD) results further suggest that the morphological diversity among BalCP20-P3 and its mutations lays on the proportion of *α*-helix. The above results demonstrate that cysteines and disulfide bonds play a crucial role in the self-assembly of BalCP20-P3. This study provides new insights into BalCP20 underwater adhesion, and brings in new inspirations for the development of novel bionic underwater adhesive.

## 1 Introduction

Biofouling occurs on all marine facilities surfaces by microorganisms, animals and plants, which either increases ship resistance, fuel consumption or severe metal corrosions, causing great economic costs [1; 2; 3]. Among those fouling organisms, barnacles are renowned for their ability to remain attached throughout their adult life [4]. Barnacles strongly adhere to immersed solid substrates using a mixture of cement proteins (CPs) that self-assembles into a permanently bonded layer [5]. Their base plates consist of cement protein will keep binding to foreign surfaces even when they are dead and lost all soft bodies. Hence, on the purpose of developing new technology for either antifouling or underwater adhesion, it is important to comprehensively explore the molecular mechanism of barnacle cement proteins.

So far, six CPs have been characterized and identified by their apparent molecular weighs: CP100, CP68, CP52, CP20, CP19, CP16 [6; 7; 8; 9]. These cement proteins have different spatial distribution and diverse functions. For example, the insoluble proteins CP100 and CP52 may provide bulk properties in the barnacle cement, CP19, CP20 and CP68 have been speculate to be interfacial proteins and associate with surface functions, whereas CP16 is a minor constituent and shares homology with a lysozyme-like enzyme, which is proposed to remove biofilm from the substratum and/or protect the cement from microbial degradation [10]. Moreover, CP20 is also found to have adsorption activity to calcite, indicating its dual function for surface binding and biomineralization [8].

Kamino suggested that the high abundant Cys residues in *Megabalanus rosa* CP20 resulted in the formation of intramolecular disulfide bonds, which are essential for the proper folding of the monomeric protein structure [8]. NMR and molecular dynamics (MD) simulation results shown that rMrCP20 contains three main folded domain regions intervened by two dynamic loops, resulting in multiple protein conformations that exist in equilibrium. Besides, 12 out of 32 Cys in the rMrCP20 sequence engage in the formation of disulfide bonds [11]. Kumar also used MD simulations to investigate the molecular interactions between rMrCP20 and calcium carbonate and reported Ca^2+^ and CO_3_^2-^ ions were sequestered by protein charged surfaces [5]. More recently, breakthroughs have been made by Ali Miserez group. They successfully observed CaCO_3_ mineralization pathway regulated by rMrCP20, as well as the adhesive nanofibrils formed by its self-assembling. However, most researches are focused on recombinant *Megabalanus rosa* CP20. CP20 from other species are rarely studied.

In our previous study, we expressed recombinant *Balanus albicostatus* CP20 in *E*.*Coli*. Though the expression yield is very high, purified protein tend to separate out from the solution and form significant flocculent precipitation after stored in 4°C (data not shown). Since there are 18 cysteines in BalCP20 sequence, intramolecular or intermolecular disulfide bonds are likely to form, as reported by Kamino and others [8; 11]. Disulfide bonds are essential for the proper folding of monomeric proteins on one hand. On the other hand, the misplace of disulfide bonds will cause protein misfolding and precipitation. Based on this consideration, we suspend our study on whole-length BalCP20, and turn to explore its derived peptides functions and properties. Through sequence comparison, one section of BalCP20 is found to share high homology with the EGF calcium binding domain mussel foot protein 2 (mfp-2). We name it as BalCP20-P3. The BalCP20-P3 peptide is likely to be the key to decipher CP20 functions and associated mechanism.

In this study, AFM and TEM investigations show that BalCP20-P3 forms grain-like spindles, which further hierarchically assembly into fractal-like structures looks like ears of wheat. SDS-PAGE, AFM and LSCM show that DTT treatment opens up disulfide bonds between cysteines and disrupts the fractal-like structures (eras of wheat). To validify the essential role of cysteines and disulfide bonds for CP20 self-assembly, four mutants of BalCP20-P3 (C→A) are designed. It is significant that the grain-like spindles and ears-of-wheat-like structures are abolished when one of the BalCP20-P3 four cysteines is mutated by alanine. Circular dichroism (CD) results further indicate that the secondary structure proportion of BalCP20 is substantially different. The unique morphology of BalCP20-P3 lays on the abundance of *α*-helix. The above results demonstrate that cysteines and disulfide bonds play a crucial role in the self-assembly of BalCP20-P3. This study provides new insights into BalCP20 underwater adhesion, and brings in new inspirations for the development of novel bionic underwater adhesive.

## 2 Materials and Methods

### 2.1 Materials

Peptides are chemically synthesized, Milli-Q water (Milli-Q system, Millipore, Bedford, MA), Dithiothreitol (DTT); Thioflavin T (ThT) (Sangon Bioengineering (Shanghai) Co., Ltd. Tricine SDS Sample Buffer; SeeBlueTM Plus2 Prestained Standard; NovexTM 10-20% Tricine Gel, Tricine SDS Running Buffer (10×) (ThermoFisher SCIENTIFIC, USA).

### 2.2 Preparation of Peptides

The peptide designed in this study is derived from the third repeat unit in the primary structure of BlaCP20 (amino acid sequence see ref. [12]). The full-length amino acid sequence of BalCP20 was obtained from the NCBI (National Center for Biotechnology Information) database, and divided into 4 repeats according to the conserved cysteines (Fig. 1A). The third repeat peptide (P3) of BalCP20 has high similarity with the epidermal growth factor (EGF) domain of mussel foot protein 2 (mfp-2), especially the locations of conserved cysteines (Fig. 1B). The conserved cysteines in the EGF calcium binding domain (EGF-CBD) helps mfp-2 maintain correct tertiary structure by forming intramolecular disulfide bonds [13]. In this point, the P3 sequence is likely to be the functional unit of BalCP20. To further investigate the effect of the number and position of cysteines, series mutants of BalCP20-P3 are designed by alanine scanning. Designed peptides are synthesized by Fmoc solid-phase [14] and purified by reverse-phase high-performance liquid chromatography using a Zorbax 300SB-C18 column (4.6mm×150mm; Agilent Technologies, Palo Alto, CA). Elution was conducted with a water-acetonitrile liner gradient (100% (v/v) acetonitrile) containing 0.1% (v/v) trifluoroacetic acid. The molecular mass of the purified peptide was confirmed by liquid chromatography/mass spectroscopy using a Water 2695 HPLC Separations Module (Waters Alliance, USA). (S1-S5). The synthesized peptides were dissolved in ultrapure water (Milli-Q system, Millipore, Bedford, MA), store at 4°C for later use.

**Figure 1.**
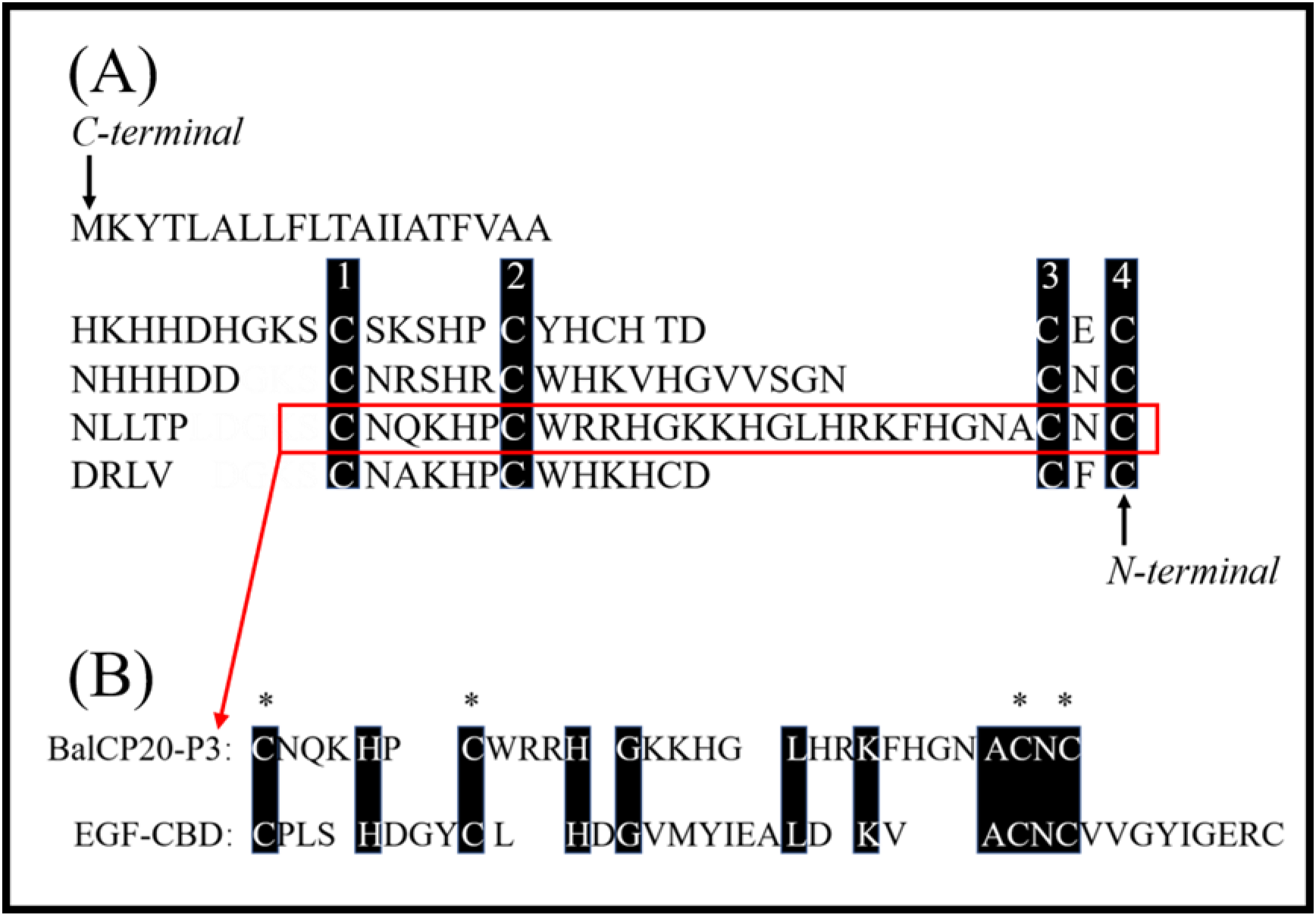
Representation of the repetitive unit of BalCP20 by the alignment of Cys residues and sequence comparison of BalCP20-P3 and EGF-CBD. The alignment of Cys residues indicates four repetitions in the BalCP20 sequence. All the Cys residues are boxed in black. BalCP20-P3 was boxed in red. (A). The amino acid alignment of BalCP20-P3 and EGF-CBD in the calcium ion binding domain of mussel adhesion protein mfp-2 (B), in which the black boxes indicate the same amino acid residues, and * indicates that the two sequences are conserved and identical of cysteine.

### 2.3 Visualization with SEM, TEM, AFM and LSCM

Before SEM analysis, peptides powders were dissolved in Milli-Q at 4 °C. 10 μl of BalCP20-P3 peptide solution was then placed on a clean silicon wafer and briefly dried at ambient temperature. Samples on silicon wafer were then observed an S-4800 scanning electron microscope (Hitachi, Japan).

For the AFM analysis, a 10 μl of peptide Milli-Q solution was placed on the newly peeled surface of mica wafer and briefly dried at ambient temperature. AFM images was captured in the tapping mode by a Nano Scope-ScanAsyst in air. wks controller (Bruker Corporation, DE) and intelligent scanning (ScanAsyst in AIR), using a silicon probe with a cantilever length of 115 μm and a spring constant of 0.4 N/m. Images with a scanning range of 0.6-8 μm were taken at a scanning rate of 1 Hz with 256 lines per image. LSCM was photographed with TCS-SP8 (leica MICROSYSTEMS, DE), and the photographed results were processed with ImageJ software.

For the transmission electron microscopy (TEM) analysis, 10 μl of peptide Milli-Q solution was placed on a 200-mesh copper mesh support film (Beijing Zhongjingkeyi Technology Co., Ltd.). The copper mesh was kept wet for 5 min, then 10 μl aliquot of uranyl acetate (mass fraction 2%, Hubei Chushengwei Chemical Co., Ltd.) was stained for 5 minutes and dried at room temperature. TEM images were recorded using a transmission electron microscope TEM (HT-7700, Hitachi, Japan) [15].

### 2.4 DTT Treatment and SDS-PAGE

SDS-PAGE was performed according to the method of Gao et al. [16] with some modifications in the electrophoresis apparatus and conditions: 10 μl of BalCP20-P3 sample prepared as described above was mixed with an equal volume of gradient DTT (4∼200 mM) and 20 μl Tricine SDS Sample Buffer (2×) (Thermo SCIENTIFIC, USA) and placed in a water bath at 70°C for 10 min. After, samples were electrophoresed with NovexTM 10-20% Tricine Gel (Thermo SCIENTIFIC, USA) at 200 V, 120 mA for 10 min and 200 V, 80 mA for 30 min and stained with Coomassie brilliant blue G-250 for 5 h, decolorize with 10% acetic acid solution.

### 2.5 Circular Dichroism (CD) and Thioflavin T (ThT)

CD data were recorded on a BRIGHTTIME Chirascan, JASCO810, Jasco-815 spectrometer, measuring from 260 nm to 190 nm at 25°C [15]. Spectra Manager software was used to smooth the CD data. Data was then saved and uploaded it to http://dichroweb.cryst.bbk.ac.uk. Wavenumber range from 190 to 260 nm were selected for calculation to obtain the relative content of secondary structure [17]. Analyses were performed as three independent samples and data were averaged.

To determine whether CP20-P3 underwent an Amyloid-like self-assembly, a Thioflavin T (ThT) test was conducted [18]. A 20 mM ThT aqueous concentrate was diluted 1000-fold with 50 mM potassium phosphate buffer (pH 6.0) immediately before being used. Aliquots of the peptide concentrate (0.2∼4 mg/mL) was mixed with equal volume of diluted ThT solution, the mixture was then incubated for 30 min at 4°C. Fluorescence was measured with a FLUOROSKAN ASCENT FL spectrometer (Thermo SCIENTIFIC, USA) at 25°C with an excitation wavelength of 430 nm and an emission wavelength of 481 nm. The fluorescence intensity of the blank solution without peptide was subtracted as background. Peptides with fluorescence intensities of > 0.5 were defined as “ThT binding” in the study [17].

## 3 Results

### 3.1 Morphological observation by SEM/TEM/AFM

Morphological observations of BalCP20-P3 dissolved in Milli-Q are presented in Fig. 2. SEM analysis reveals the formation of compact stacked peptide pipes within hours after the BalCP20-P3 were dissolved in Milli-Q. The peptide-pipes are of diameters around one micron, and almost no amorphous structures were observed within the entire sample (Fig. 2A, B). In order to get further insights regarding the structural characteristics of the stacked peptides, samples are analyzed by high-resolution AFM and TEM. Assembled peptide structures deposited on mica further formed organized and regular “wheat spike-like” structures with a three-dimensional conformation. AFM analysis clearly indicated that the wheat spike-like structures are about 1 μm in diameter, while the length and width of their subunits looks like wheat grains are about microns and hundreds of nanometers (Fig. 2C, D). Moreover, TEM analysis showed that the peptide of BalCP20-P3 assemblies into short spindles about 100∼200 nm, which cross-linked to each other end-to-end (Fig. 2E, F). It is speculated that the differences during sample preparation results in the diversity of peptide morphology. For TEM, samples were dropped on copper meshes. During the desiccation at room temperature, peptides sink into micropores will not influenced by the changes of aqueous tension. However, for SEM, there will be a liquid flow acts on peptide self-assemblies during desiccation, leading to the sample gathering in certain regions and the formation of larger grains and even wheat spikes. As to the difference between SEM and AFM results, the vacuum operation conditions of SEM is likely to contributes to the transition from wheat spike into stacked pipes.

**Figure 2.**
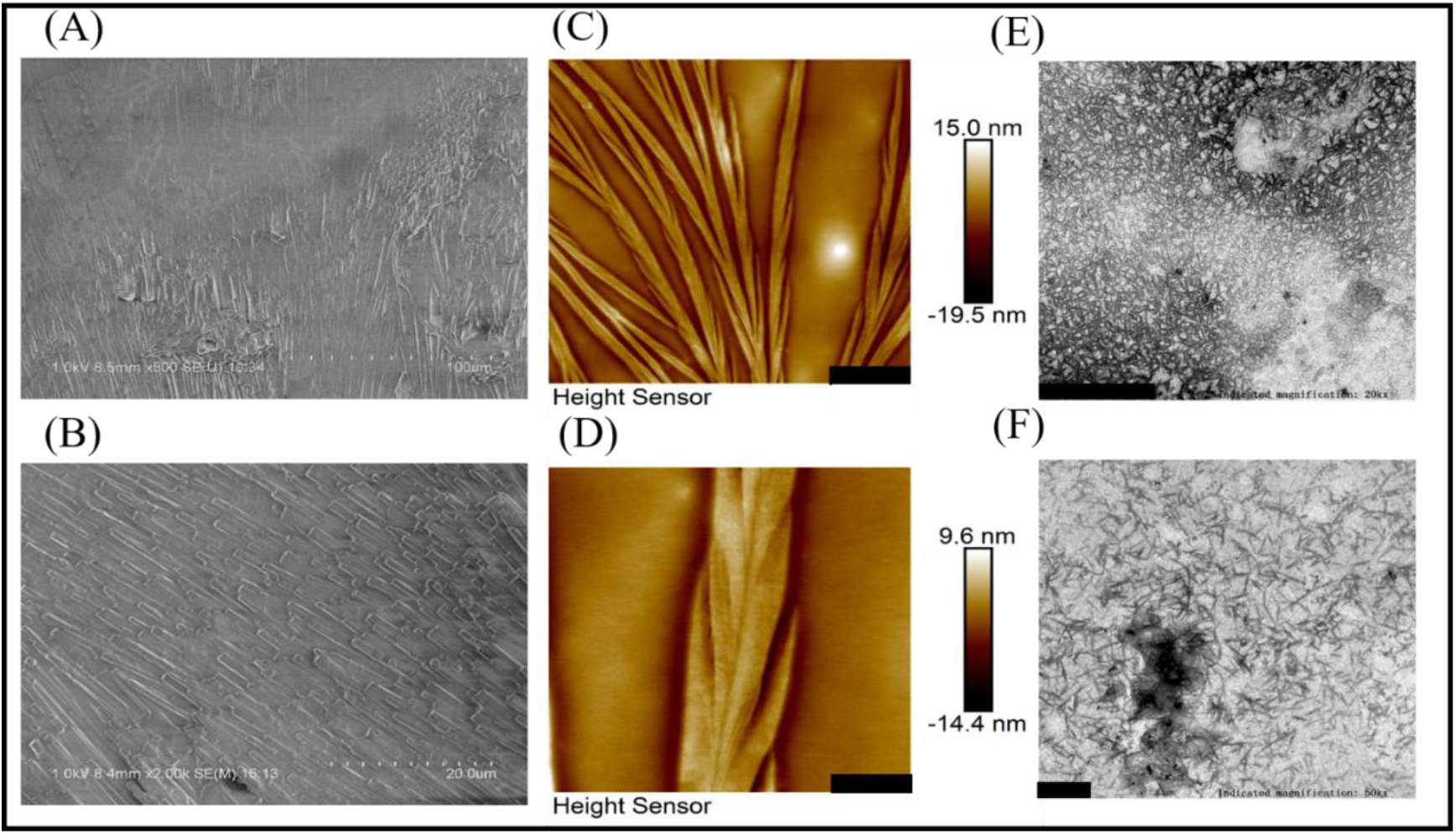
Microscopic characterization of BalCP20-P3. SEM of BalCP20-P3. Scale bar = 100 μm (A). Scale bar = 20 μm (B). AFM of BalCP20-P3. Scale bar = 3 μm (C). Scale bar = 1 μm (D). TEM of BalCP20-P3. Scale bar = 1 μm (E). Scale bar = 200 μm (F).

### 3.2 SDS-PAGE/AFM/LSCM of BalCP20-P3 after DTT treatment

BalCP20-P3 solutions were treated with series concentrations of DTT and evaluated by SDS-PAGE (Fig. 3A). For BalCP20-P3 treated with low DTT concentrations or no DTT, there are obvious smears in the lands. As the DTT concentrations goes up, smears gradually disappears and the bands tend to enrich around 3 kDa. This indicates that the disulfide bonds among BalCP20-P3 were disrupted, and oligomer or monomers were released. Correspondingly, the grain-like or wheat spike-like morphology of BalCP20-P3 also disappears. Both LSCM and AFM detected spherical structures in DTT-treated samples. Since these spherical particles were flowing in the droplets, it is difficult for LSCM to determine a focal plane (Fig3.B). AFM accurately described the spherical particles with diameters approximately 150 nm (Fig 3C, D, E), and evenly distributed in the entire area.

**Figure 3.**
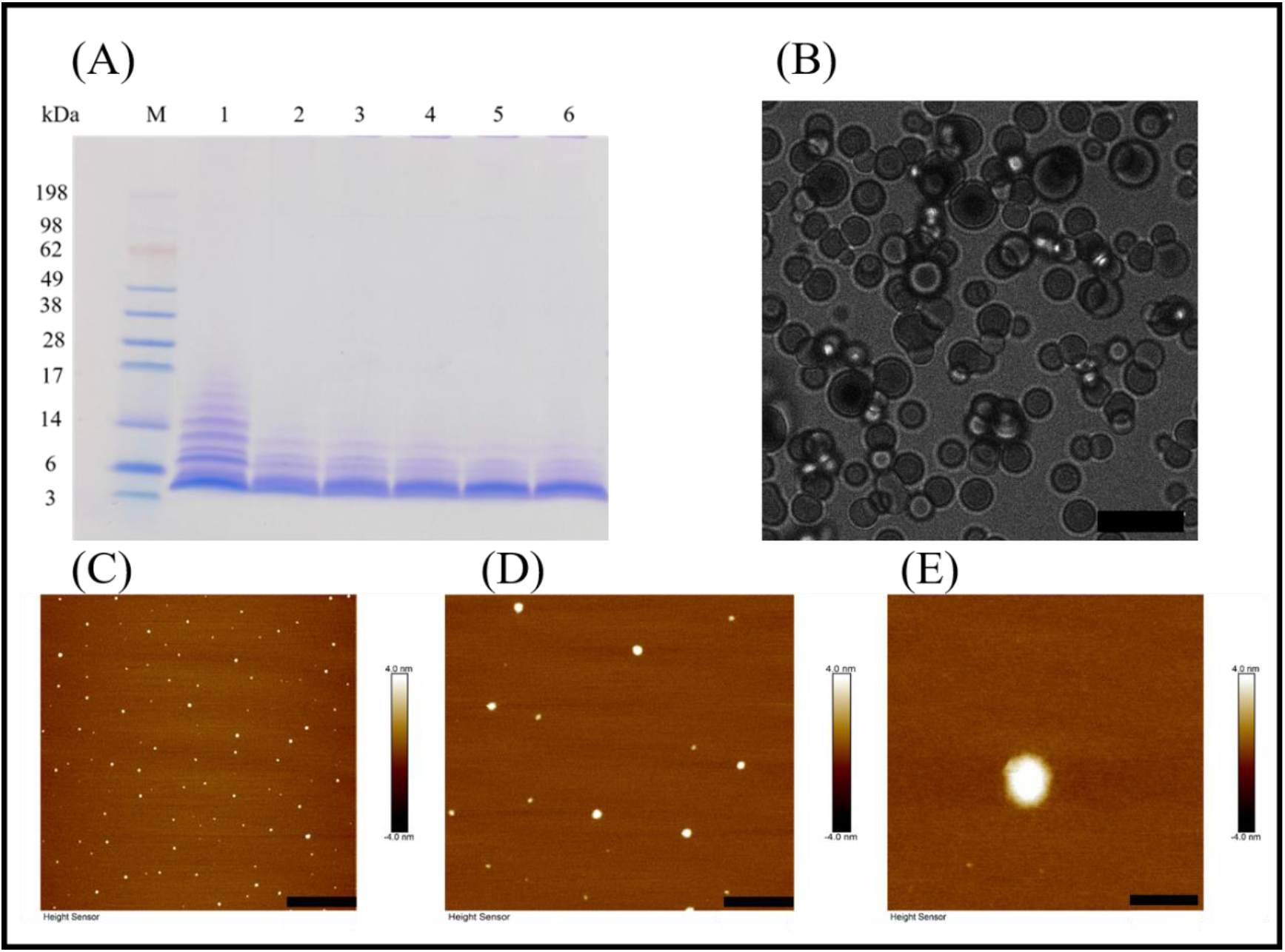
BalCP20-P3 after DTT treatment. SDS-PAGE of BalCP20-P3±DTT (A). M is Mark protein. Lane 1 is BalCP20-P3 of -DTT, the distribution of protein bands ranges from 3 to 17 kDa, and BalCP20-P3 forms multiple multimers. Lane 2 is BalCP20-P3 treated by 1 mM DTT, the multimer gradually dissociates and the molecular wight was between 3 and 15 kDa. Lane 3 is BalCP20-P3 treated by 5 mM DTT and Lane 4 is BalCP20-P3 treated by 10 mM DTT. The protein bands of the two lanes were concentrated in 3 to 14 kDa, and the band near 3 kDa is more obvious. 30 mM and 50 mM DTT was added to lane 5 and lane 6, respectively, the protein bands were mainly around 3 kDa. (BalCP20-P3 _MW_ = 3.35389 kDa). LSCM of BalCP20-P3 after DTT treatment (B). BalCP20-P3 exhibits a regular spherical shape. Scale bar = 10 μm. AFM of BalCP20-P3 after DTT treatment. Spherical particles evenly distributed in the field of view. Scale bar = 3 μm (C). Scale bar = 1 μm (D). Scale bar = 200 nm (E).

### 3.3 AFM/CD/ThT of BalCP20-P3 mutants

To verify the fundamental role of cysteines or disulfide bonds, four mutants were designed by change one of the four cysteines in BalCP-P3 to alanine (Fig. 4A). Firstly, the different self-assembly properties among BalCP20-P3 and mutants were observed by AFM. For wildtype BalCP20-P3, it is a delicate wheat spike hierarchical structure. However, these morphological features are entirely abolished when merely one cysteine was mutated. AFM observed spheres with diameters of approximate 500 nm in BalCP20-P3-M1 and BalCP20-P3-M4 samples (Fig. 4C, F), while short rods with length about 1 micron in BalCP20-P3-M2∼M4 (Fig. 4 D, E, F). Furthermore, circular dichroism spectrum was conducted to explain their different self-assembly behaviors at the level of secondary structure. Unordered and turn structures typically show negative peaks near 195 nm, while the *β*-sheet generally shows positive near 195 nm and negative peaks near 220 nm. *α*-helix structures typically show positive peaks near 190 nm and a negative peak at 222 nm [19]. The proportions of secondary structures of BalCP20-P3 and its mutants are presented in Table 1. The secondary structure of BalCP20-P3 is dominated by *α*-helix. The secondary structures of the mutants of BalCP20-P3 are dominated by *β*-sheet and unordered. Normalized root means square deviations (NRMSD) of BalCP20-P3 and its mutants below 0.15 generally indicate acceptable fits [20].

**Figure 4.**
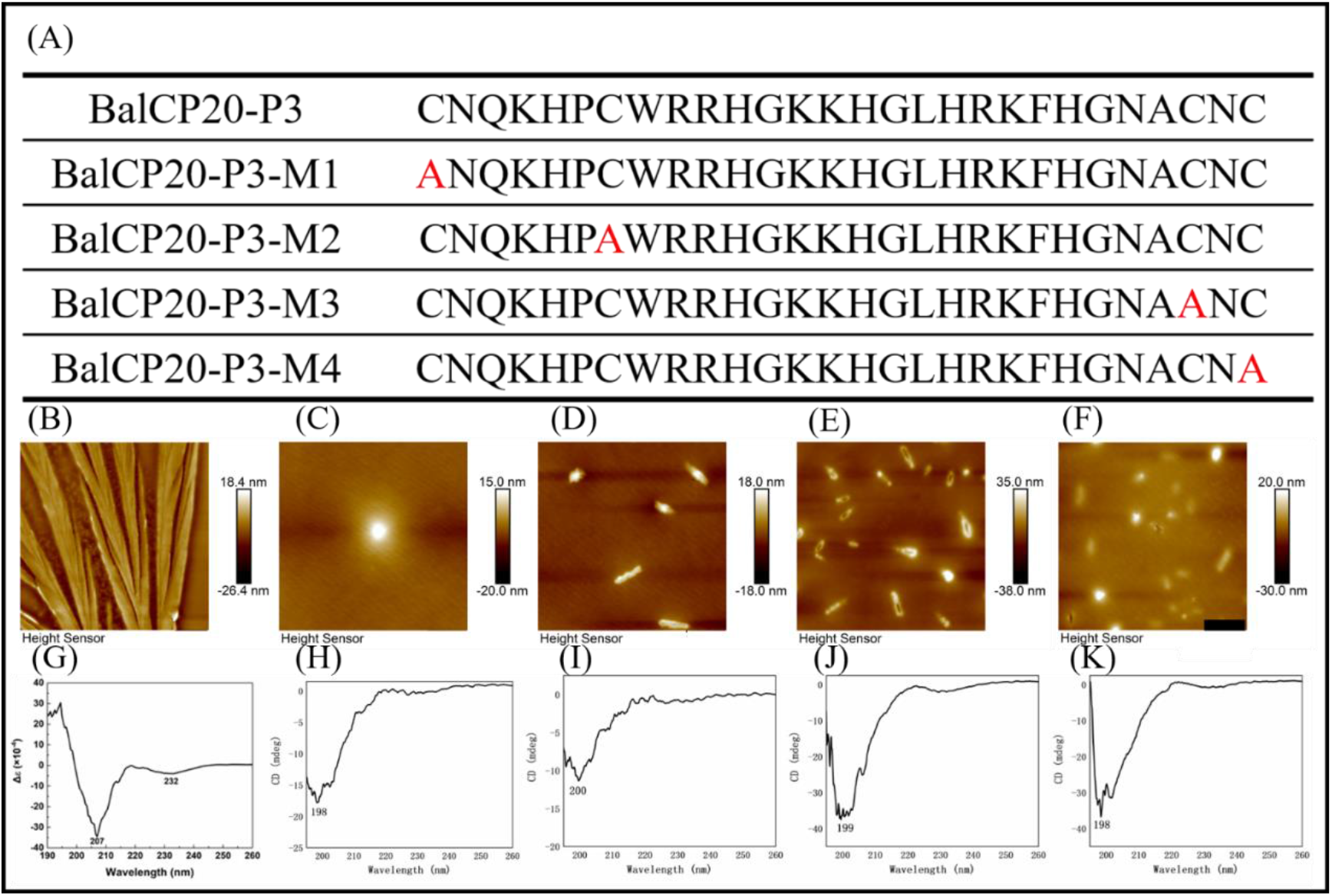
Sequence design, AFM, CD of BalCP20-P3 and its mutants. BalCP20-P3 and its mutant amino acid sequence design (A). Mutants mutated amino acid residues were marked in red. BalCP20-P3 presents a wheat spike-like structure (B). Self-assembly of BalCP20-P3-M1 was spherical (C). BalCP20-P3-M2 (D) and BalCP20-P3-M3 (E) were a rod-like structure of varying lengths. BalCP20-P3-M4 has either spherical or rod-like structures (F). The scale bar of all AFM images was 1 μm. CD of BalCP20-P3 (G). CD of BalCP20-P3-M1 (I). CD of BalCP20-P3-M2 (J). CD of BalCP20-P3-M3 (K). CD of BalCP20-P3-M4 (L). The abscissa is the wavelength (190 nm to 260 nm) and the ordinate is the absorbance.

**Table 1.** Secondary structure ratio of BalCP20-P3 and its mutants.

| Sample | α-helix | $3_{10}$-α-helix | β-sheet | β-turn | PP2 | unordered | NRMSD |
| --- | --- | --- | --- | --- | --- | --- | --- |
| BalCP20-P3 | 0.348 | 0.170 | 0.020 | 0.122 | 0.196 | 0.129 | *0.022* |
| BalCP20-P3-M1 | 0.004 | 0.033 | 0.144 | 0.153 | 0.165 | 0.501 | *0.072* |
| BalCP20-P3-M2 | 0.003 | 0.040 | 0.216 | 0.144 | 0.139 | 0.458 | *0.099* |
| BalCP20-P3-M3 | 0.002 | 0.032 | 0.279 | 0.115 | 0.130 | 0.442 | *0.145* |
| BalCP20-P3-M4 | 0.005 | 0.111 | 0.045 | 0.181 | 0.151 | 0.507 | *0.139* |

The normalized root mean square deviations (NRMSD) are shown in italics. NRMSD below 0.15 indicates a credible result.

## 4 Discussion

In this study, a novel peptide derived from BalCP20 amino acid sequence was found and studied. This peptide presents distinct self-assembly properties. Firstly, peptides assembly into spindles with length about hundreds of nanometers. Subsequently, these spindles further aggregated into grain-like structures and even a wheat-spike-like hierarchical morphology. Thirdly, this unique self-assembly of BalCP20-P3 is then found to related to its underwater adhesion ability.

At first place, we speculated that BalCP20 and BalCP20-P3 may underwent a amyloid-like fibrils formation process, like CP19 [21] and CP52 [17]. However, no significant fluorescence was detected by Thioflavin T tests. Therefore, CD was employed to decipher the self-assembly mechanism of BalCP20-P3. Self-assembly is a spontaneous organization that simple molecular building blocks into supramolecular structures. Typical noncovalent interactions such as hydrogen bonding, electrostatic attraction, van der walls forces and aromatic contacts will resulted this process and endows peptides with distinct secondary structures [22]. Among them, *α*-helix and *β*-sheet are two most abundant structural motifs regulate the interfacial interactions between neighboring proteins [23]. In helical peptides, the propagation of H-bonding occurs in the axial direction within the helical axis and every N-H amide proton forms H-bonding with the amide C=O group located at the fifth position. In addition, a second type of prevalent intermolecular H-bonding observed in proteins involves amino acid residues located at every first and fourth position and is known as the 3_10_-*α*-helix [24]. As circular dichroism spectrum detected, there are 34.8% *α*-helix and 17% 3_10_-*α*-helix in BalCP20-P3. However, the proportion of helix in BalCP20-P3 mutants is very rare (3∼10%). This explains why the morphology of BalCP-P3 and its mutations is so different as presented in Fig. 4. Since there is a high content of helix, BalCP-P3 molecules are twisted and their supramolecular structures looks like spindles. However, mutants of BalCP20-P3 assembly into spheres or short rods, with no tendency to twist (contort).

Moreover, we find that a typical seven repeats “c-d-e-f-g-h-a-b” existed in BalCP-P3 which contributed to the formation of helixes. In a typical “c-d-e-f-g-h-a-b” repeats, a, d represent hydrophobic amino acids, which provide a hydrophobic surface in the spiro-selective structure and act as oligo domains when multiple helical peptides are combined. e, g represents charged amino acids, which provide electrostatic action and promote helix formation, b, c, and f have little effect on the helical structure [25]. If there are “sticky end” at both ends of the helical peptide, such as cysteine [26], they will be connected to each other and extend in the lateral and longitudinal directions to form micron-scale self-assemblies.

In the paper, G-L-H-R-K-F-H in the BalCP20-P3 sequence is a typical heptad repeat. H, K can provide electrostatic force, and F and L provide hydrophobic interface. BalCP20-P3 with seven repeats first forms helical units under electrostatic and hydrophobic interactions. These helical units will further cross-link to each other and extend transversely and longitudinally to form grain-like structures by means of disulfide bonds, as detected by TEM. These grain-like structures will further gather into micron-scale wheat-spike-like or stacked pipes in AFM and SEM assays. For AFM sample preparation, BalCP20-P3 solutions are firstly deposited on the mica. As the water gradually evaporate at room temperature, the water flow will apply to the peptide assemblies. Due to the differences in viscosity between the air and solution [27], these spindles will clump together to form larger structures, says grain-like or wheat-spike like structures. As for SEM, since there is an additional vacuuming operation condition, the self-assemblies of BalCP20-P3 will be more compact and closely gathered as stacked piles. (Fig. 2A, B).

In summary, this study finds a potential function motif in BalCP20 amino acid sequence. Results presented here bring in new insights into the biochemical properties and wet adhesion mechanism of barnacle cement protein CP20, as well as new inspirations for the development of new type bionic adhesives. Gazing into futurity, efforts will be paid to study its self-assembly properties in conditions more similar to sea conditions (pH, ionic strength, etc.,) to better understand the wet adhesion mechanism of BalCP20, and to give better guidance for its bionic applications.

## 6 Author Contributions

Baoshan Li and Junyi Song designed and performed experiments, analyzed data, and completed the manuscript. Zonghuang Ye and Ling Zeng assisted in directing graduate students to carry out experiments. Ting Mao designed and plotted the schematic diagrams. Biru Hu designed experiments and wrote the manuscript. All authors contributed to the article and approved the submitted version.

## 8 Acknowledgments

The authors would like to thank Xianzhen Huang from Shiyanjia Lab (**www.shiyanjia.com**) for the AFM analysis and Central South University Xiangya Hospital for technical assistance.

## 1 Data Availability Statement

Data available on request from the authors.

## Reference

[1] R.L. Townsin, The ship hull fouling penalty. Biofouling 19 Suppl (2003) 9–15.

[2] M.P. Schultz, Effects of coating roughness and biofouling on ship resistance and powering. Biofouling 23 (2007) 331–341.

[3] D.J. Blackwood, C.S. Lim, S.L.M. Teo, X. Hu, and J. Pang, Macrofouling induced localized corrosion of stainless steel in Singapore seawater. Corrosion Science 129 (2017) 152–160.

[4] R. Cleverley, D. Webb, S. Middlemiss, P. Duke, A. Clare, K. Okano, C. Harwood, and N. Aldred, In Vitro Oxidative Crosslinking of Recombinant Barnacle Cyprid Cement Gland Proteins. Mar Biotechnol (NY) 23 (2021) 928–942.

[5] A. Kumar, H. Mohanram, J. Li, H. Le Ferrand, C.S. Verma, and A. Miserez, Disorder–Order Interplay of a Barnacle Cement Protein Triggered by Interactions with Calcium and Carbonate Ions: A Molecular Dynamics Study. Chemistry of Materials 32 (2020) 8845–8859.

[6] K. Kamino, Barnacle Underwater Attachment, Biological Adhesives, 2016, pp. 153–176.

[7] K. Kamino, M. Nakano, and S. Kanai, Significance of the conformation of building blocks in curing of barnacle underwater adhesive. FEBS J 279 (2012) 1750–1760.

[8] Y. Mori, Y. Urushida, M. Nakano, S. Uchiyama, and K. Kamino, Calcite-specific coupling protein in barnacle underwater cement. FEBS J 274 (2007) 6436–6446.

[9] Y. Urushida, M. Nakano, S. Matsuda, N. Inoue, S. Kanai, N. Kitamura, T. Nishino, and K. Kamino, Identification and functional characterization of a novel barnacle cement protein. FEBS J 274 (2007) 4336–4346.

[10] M.A. Tilbury, S. McCarthy, M. Domagalska, T. Ederth, A.M. Power, and J.G. Wall, The expression and characterization of recombinant cp19k barnacle cement protein from Pollicipes pollicipes. Philos Trans R Soc Lond B Biol Sci 374 (2019) 20190205.

[11] H. Mohanram, A. Kumar, C.S. Verma, K. Pervushin, and A. Miserez, Three-dimensional structure of Megabalanus rosa Cement Protein 20 revealed by multi-dimensional NMR and molecular dynamics simulations. Philos Trans R Soc Lond B Biol Sci 374 (2019) 20190198.

[12] L.S. He, G. Zhang, and P.Y. Qian, Characterization of two 20kDa-cement protein (cp20k) homologues in Amphibalanus amphitrite. PLoS One 8 (2013) e64130.

[13] D.S. Hwang, and J.H. Waite, Three intrinsically unstructured mussel adhesive proteins, mfp-1, mfp-2, and mfp-3: analysis by circular dichroism. Protein Sci 21 (2012) 1689–1695.

[14] M.E. Kim, J.K. Seon, J.Y. Kang, T.R. Yoon, J.S. Lee, and H.K. Kim, Bone-Forming Peptide-4 Induces Osteogenic Differentiation and VEGF Expression on Multipotent Bone Marrow Stromal Cells. Front Bioeng Biotechnol 9 (2021) 929.

[15] S. Sonzini, S.T. Jones, Z. Walsh, and O.A. Scherman, Simple fluorinated moiety insertion on Abeta 16-23 peptide for stain-free TEM imaging. Analyst 140 (2015) 2735–2740.

[16] L. Gao, W. Ma, J. Chen, K. Wang, J. Li, S. Wang, F. Bekes, R. Appels, and Y. Yan, Characterization and comparative analysis of wheat high molecular weight glutenin subunits by SDS-PAGE, RP-HPLC, HPCE, and MALDI-TOF-MS. J Agric Food Chem 58 (2010) 2777–2786.

[17] M. Nakano, and K. Kamino, Amyloid-like conformation and interaction for the self-assembly in barnacle underwater cement. Biochemistry 54 (2015) 826–835.

[18] A.B. Matiiv, N.P. Trubitsina, A.G. Matveenko, Y.A. Barbitoff, G.A. Zhouravleva, and S.A. Bondarev, Amyloid and Amyloid-Like Aggregates: Diversity and the Term Crisis. Biochemistry (Mosc) 85 (2020) 1011–1034.

[19] D.E. Barlow, G.H. Dickinson, B. Orihuela, J.L. Kulp, 3rd, D. Rittschof, and K.J. Wahl, Characterization of the adhesive plaque of the barnacle Balanus amphitrite: amyloid-like nanofibrils are a major component. Langmuir 26 (2010) 6549–6556.

[20] B.R. Baker, and R.L. Garrell, G-factor analysis of protein secondary structure in solutions and thin films. Faraday Discuss 126 (2004) 209–22; discussion 245-254.

[21] X. Liu, C. Liang, X. Zhang, J. Li, J. Huang, L. Zeng, Z. Ye, B. Hu, and W. Wu, Amyloid fibril aggregation: An insight into the underwater adhesion of barnacle cement. Biochem Biophys Res Commun 493 (2017) 654–659.

[22] T. Aida, E.W. Meijer, and S.I. Stupp, Functional supramolecular polymers. Science 335 (2012) 813–817.

[23] J.S. Richardson, The anatomy and taxonomy of protein structure. Advances in protein chemistry 34 (1981) 167–339.

[24] S. Mondal, and E. Gazit, The Self-Assembly of Helical Peptide Building Blocks. ChemNanoMat 2 (2016) 323–332.

[25] S.a. Potekhin, T. Melnik, V. Popov, N. Lanina, A.a. Vazina, P. Rigler, A. Verdini, G. Corradin, and A. Kajava, De novo design of fibrils made of short α-helical coiled coil peptides. Chemistry & biology 8 (2001) 1025–1032.

[26] M. Depuydt, J. Messens, and J.-F. Collet, How proteins form disulfide bonds. Antioxidants & redox signaling 15 (2011) 49–66.

[27] J. Chen, S. Qin, X. Wu, and A.P. Chu, Morphology and Pattern Control of Diphenylalanine Self-Assembly via Evaporative Dewetting. ACS Nano 10 (2016) 832–838.

