## Supplementary Figures 1-5 for "The essential role of disulfide bonds for the hierarchical self-assembly and wet-adhesion of CP20-derived peptides"

### *Supplementary Material*

#### 1.1 Supplementary Figures

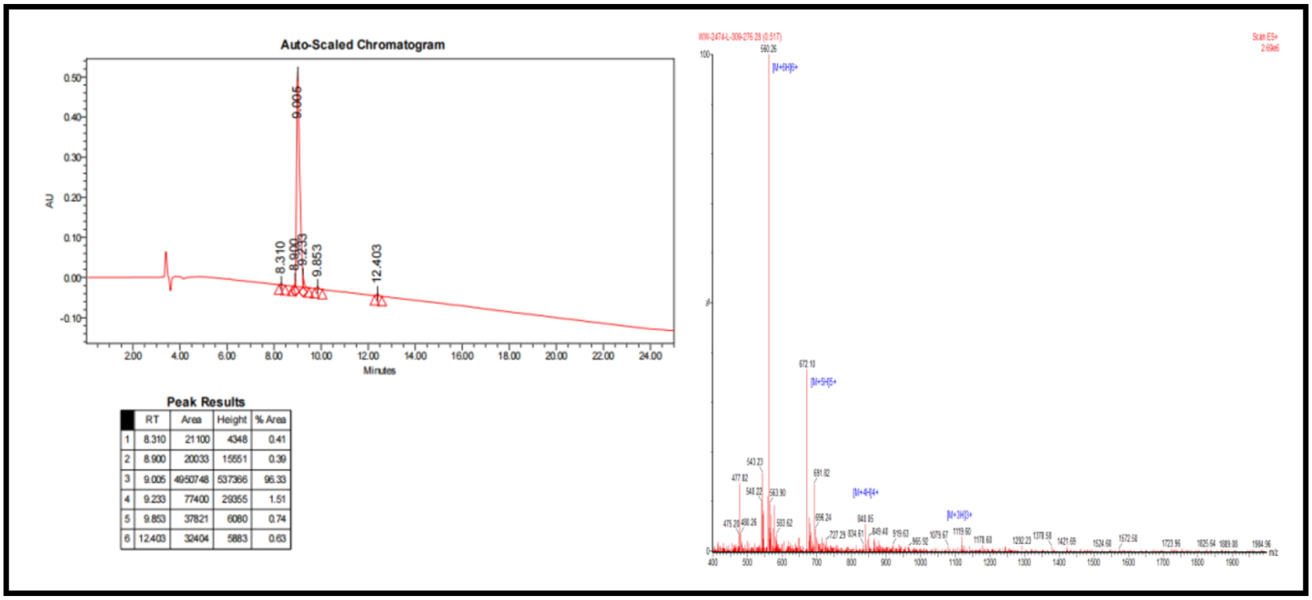


**Supplementary Figure 1.** HPLC of BalCP20-P3 (left), MS of BalCP20-P3 (right)

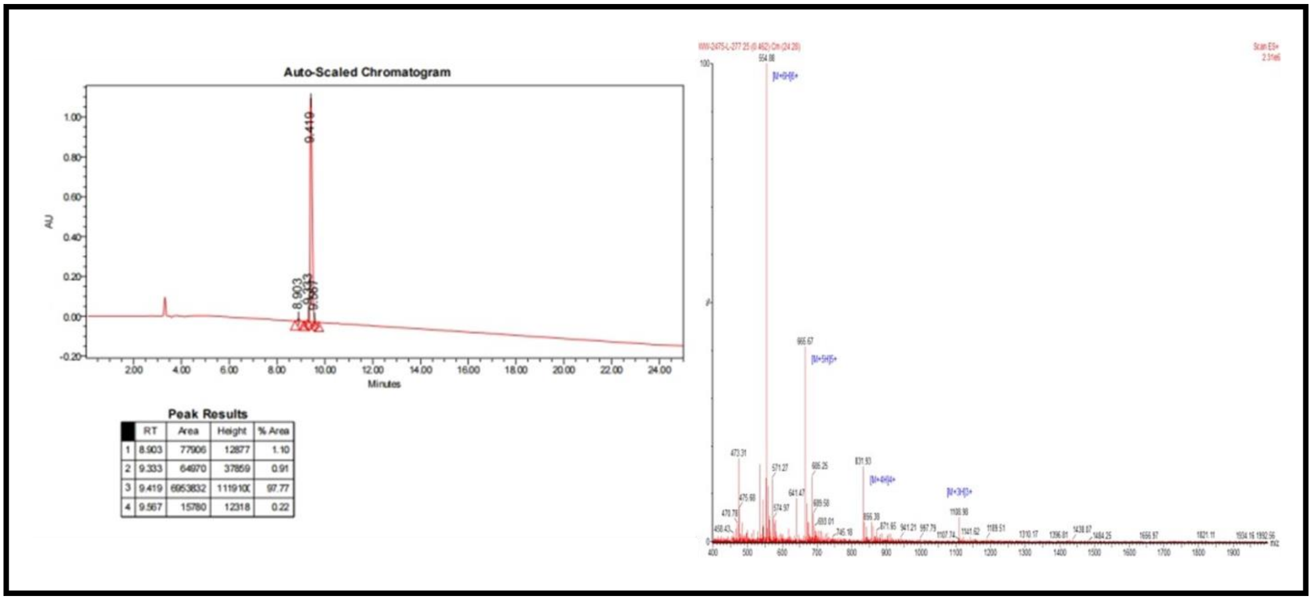


**Supplementary Figure 2.** HPLC of BalCP20-P3-M1 (left), MS of BalCP20-P3-M1 (right)

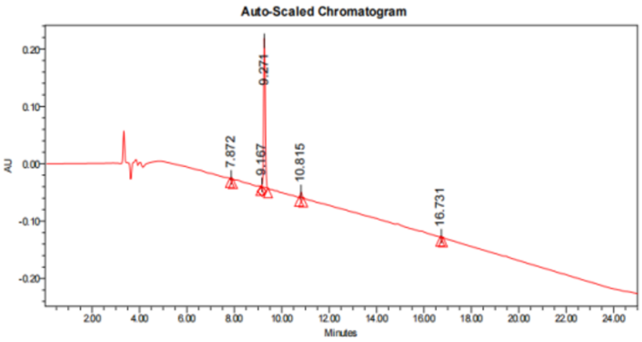


**Peak Results**

| | RT | Area | Height | % Area |
|---|---|---|---|---|
| 1 | 7.872 | 8468 | 2132 | 0.66 |
| 2 | 9.167 | 5852 | 3246 | 0.45 |
| 3 | 9.271 | 1222886 | 257051 | 95.07 |
| 4 | 10.815 | 42537 | 9133 | 3.31 |
| 5 | 16.731 | 6569 | 1566 | 0.51 |

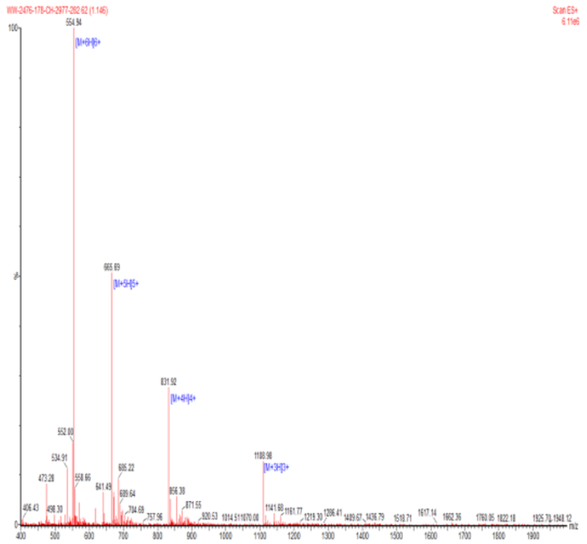

**Supplementary Figure 3.** HPLC of BalCP20-P3-M2 (left), MS of BalCP20-P3-M2 (right)

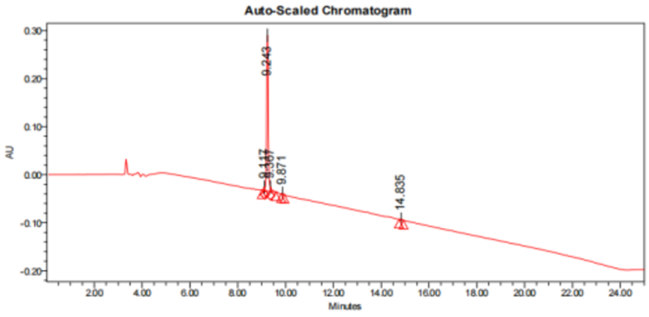


**Peak Results**

| | RT | Area | Height | % Area |
|---|---|---|---|---|
| 1 | 9.117 | 25140 | 7236 | 1.62 |
| 2 | 9.243 | 1477617 | 323297 | 95.23 |
| 3 | 9.367 | 33661 | 8930 | 2.17 |
| 4 | 9.871 | 7273 | 1734 | 0.47 |
| 5 | 14.836 | 7941 | 1586 | 0.51 |

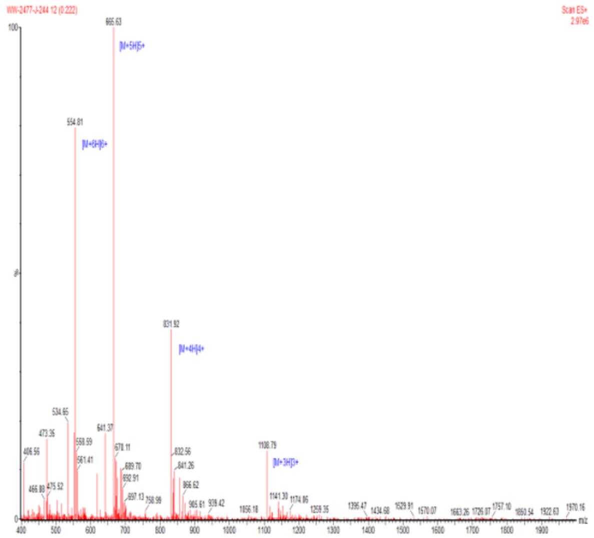

**Supplementary Figure 4.** HPLC of BalCP20-P3-M3 (left), MS of BalCP20-P3-M3 (right)

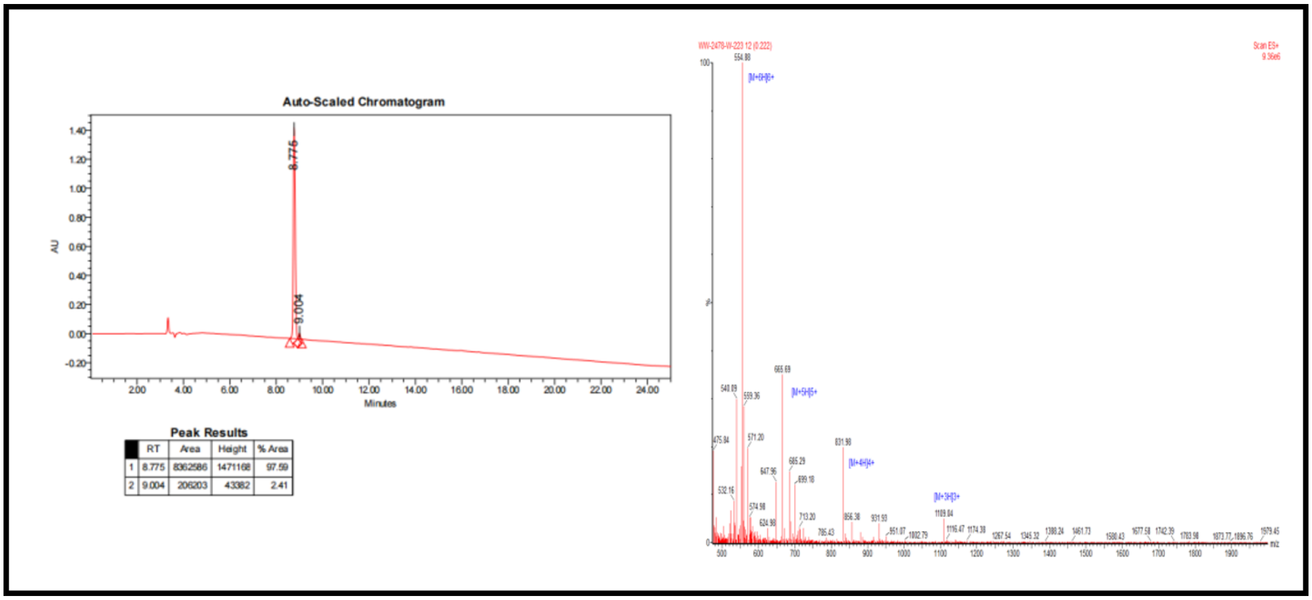


Peak Results

| | RT | Area | Height | % Area |
|---|---|---|---|---|
| 1 | 8.775 | 8362586 | 1471168 | 97.59 |
| 2 | 9.004 | 206203 | 43382 | 2.41 |

**Supplementary Figure 5.** HPLC of BalCP20-P3-M4 (left), MS of BalCP20-P3-M4 (right)
